## Supplementary Materials and Results for "Effects of early life adversity on immediate early gene expression: systematic review and 3-level meta-analysis of rodent studies"

### 1. Supplementary Methods

#### 1.1 Study Protocol

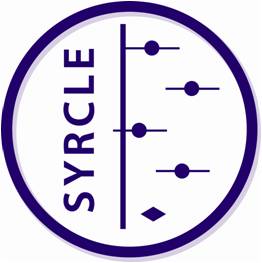

| **Systematic Review Protocol for Animal Intervention Studies**  **Format by SYRCLE (**[**www.syrcle.nl**](http://www.syrcle.nl)**)**  **Version 2.0 (December 2014)** | | | |
| --- | --- | --- | --- |
| **Item #** | **Section/Subsection/Item** | **Description** | **Changes from original protocol** |
|  | A. General | | |
| 1. | Title of the review | Effect of Early Life Adversity on Immediate Early Gene Expression in Rodents |  |
| 2. | Authors (names, affiliations, contributions) | Valeria Bonapersona^1^, Heike Schuler^1^, Marian Joëls^1,2^, R. Angela Sarabdjitsingh^1^  ^1^ Department of Translational Neuroscience, UMC Utrecht Brain Center, University Medical Center Utrecht, Utrecht University, The Netherlands  ^2^ University Medical Center Groningen, University of Groningen, The Netherlands | The protocol was written in April 2019 and since then unaltered. |
| 3. | Other contributors (names, affiliations, contributions) | SYRCLE (SYstematic Review Center for Laboratory animal Experimentation), Radboud University Nijmegen Medical Center, Nijmegen, The Netherlands |  |
| 4. | Contact person + e-mail address | Valeria Bonapersona; |  |
| 5. | Funding sources/sponsors | The Consortium on Individual Development (CID) is funded through the Gravitation program of the Dutch Ministry of Education, Culture, and Science and the Netherlands Organization for Scientific Research (NWO grant number 024.001.003). |  |
| 6. | Conflicts of interest | None |  |
| 7. | Date and location of protocol registration | Date:  Location: [www.crd.york.ac.uk](http://www.crd.york.ac.uk) |  |
| 8. | Registration number (if applicable) |  |  |
| 9. | Stage of review at time of registration | Completed preliminary searches, started with piloting of the study selection process. |  |
|  | B. Objectives | | |
|  | Background | | |
| 10. | What is already known about this disease/model/intervention? Why is it important to do this review? | Exposure to adversities during childhood (early-life adversities, ELA) increases the risk to develop psychiatric disorders in adulthood. Building upon the compelling epidemiological evidence, rodent studies have investigated the mechanistic effects of ELA on the brain, ultimately leading to changes in behavior. Modelled as alterations in maternal care, ELA alters brain development on multiple levels, including synaptic organization.  Important contributors to synaptic development and cognition are immediate early genes (IEGs). IEGs are expressed directly but transiently upon cell activity; hence, they can be considered a marker for information processing in the brain. Since IEG proteins vary from transcription factors to post-translational proteins, their different functions can highlight different aspects of synaptic development.  Systematically reviewing the current literature on the topic can provide insights on long-term changes of ELA on IEGs throughout the brain, thereby providing possible mechanisms for ELA-induced changes in information processing. |  |
|  | Research question | | |
| 11. | Specify the disease/health problem of interest | Childhood maltreatment; Early life adversity; Stress-related psychopathology; Healthy animals |  |
| 12. | Specify the population/species studied | mice and rats, because they are the most frequently used animal models in stress research; female and male |  |
| 13. | Specify the intervention/exposure | 1) Early life adversity starting before P14; early life adversity defined as alteration in maternal care; models included are maternal separation/deprivation, isolation, limited bedding and nesting, licking and grooming (as measure of variation in maternal care, with final comparisons between offspring receiving low vs high maternal care (for reference: Liu et al., 1997)), handling 2) Acute stressors applied to adult animals; will be restricted to most common ones as based on results from formal screening   Both, 1) and 2) need to be applied. |  |
| 14. | Specify the control population | Control animals differ from experimental animals only by exposure to early life adversity. |  |
| 15. | Specify the outcome measures | Immediate early gene mRNA expression, as measured by fold-change or percentage compared to control.  Immediate early gene protein expression, as measured by optical density of counts, absolute counts, or optical density of Western Blots. |  |
| 16. | State your research question (based on items 11-15) | In (healthy) mice and rats, what is the effect of childhood maltreatment, early life adversity and/or stress-related psychopathology on immediate early gene mRNA expression?  **Hypothesis-confirming research Questions:**   - In mice and rats, does early life adversity alter immediate early genes expression after an acute stress challenge? - Is this differential expression amplified by multiple hits?   **Secondary exploratory research questions:**   - Do the brain regions involved in the stress response a different sensitivity with regards to acute stress as seen in immediate early gene expression? - Are the above-mentioned effects sensitive to 1) the type of acute stressor and 2) the choice of early life adversity model? - What is the relationship in mRNA and protein expressions of any given immediate early gene in response to early life adversity? | Change: due to the uneven distribution across subgroups, secondary exploratory questions became sensitivity analyses. |
|  | C. Methods | | |
|  | Search and study identification | | |
| 17. | Identify literature databases to search (*e.g.* Pubmed, Embase, Web of science) | ☑MEDLINE via PubMed □Web of Science  □SCOPUS ☑EMBASE  □Other, namely:  □Specific journal(s), namely: |  |
| 18. | Define electronic search strategies (*e.g.* use the [step by step search guide^15^](http://www.ncbi.nlm.nih.gov/pmc/articles/PMC3265183/pdf/LA-11-087.pdf) and animal search filters[^20,^](http://www.ncbi.nlm.nih.gov/pmc/articles/PMC3104815/pdf/LA-09-117.pdf) [^21^](http://lan.sagepub.com/content/48/1/88.full.pdf+html)) | When available, please add a supplementary file containing your search strategy: **[insert file name]** |  |
| 19. | Identify other sources for study identification | ☑Reference lists of included studies □Books  ☑Reference lists of relevant reviews  □Conference proceedings, namely:  □Contacting authors/ organisations, namely:  □Other, namely: |  |
| 20. | Define search strategy for these other sources | Once the second phase of screening is completed, the reference list of the included studies and relevant reviews will be checked by one reviewer (HS). Studies that fit the search criteria identified in Questions 23-30 will be included. |  |
|  | Study selection | | |
| 21. | Define screening phases (*e.g.* pre-screening based on title/abstract, full text screening, both) | 1) Title/abstract screening 2) Full text screening |  |
| 22. | Specify (a) the number of reviewers per screening phase and (b) how discrepancies will be resolved | Study inclusion is performed by (at least) two experimenters independently and it consists of two phases. During the first phase, titles and abstracts are screened and studies are excluded if: 1) not primary publication, 2) not in mice or rats, 3) not concerning early life adversity. During the second phase, the full text is screened and studies are selected according to the priority list below. Discrepancies will be resolved by discussion between two experimenters. Should no conclusion be reached between two experimenters (VB & HS), a third researcher (RAS), will be consulted for a solution. |  |
|  | *Define all inclusion and exclusion criteria based on:* | | |
| 23. | Type of study (design) | **Inclusion criteria:** primary publications  **Exclusion criteria:** reviews; unpublished data; commentaries |  |
| 24. | Type of animals/population (*e.g.* age, gender, disease model) | **Inclusion criteria:** adult mice or rats (older than 8 weeks, but younger than 1 year); female and male  **Exclusion criteria:** any other species than mice or rats; sexes are pooled; sex is not specified; ovariectomized females; specific pathogen free animals; genetic manipulations; animals bred for high/low anxiety or novelty response or sensitivity/resilience to depression; animals separated in high/low performance; any manipulations to earlier generations; animals with any comorbidities |  |
| 25. | Type of intervention (*e.g.* dosage, timing, frequency) | **Inclusion criteria:**   1. Early life adversity starting before P14; early life adversity defined as alteration in maternal care; models included are maternal separation/deprivation, isolation, limited bedding and nesting, licking and grooming (as measure of variation in maternal care, with final comparisons between offspring receiving low vs high maternal care (for reference: Liu et al., 1997)); handling is also considered early life adversity, but will be included only at a systematic review level 2. Acute stressors applied to adult animals; the types will be restricted to most common ones as based on results from formal screening   **Exclusion criteria:** pharmacological intervention (“control” injections (of any pharmacological intervention) such as vehicle, saline, sesame oil are instead included); communal nesting as early life adversity model; maternal separation with early weaning, unless early weaning is also applied to control group; the same acute stressor has been applied earlier in life and is therefore not new to the animal | Change: we include also at rest measures. I.e. lack of acute stress is not an exclusion criteria. |
| 26. | Outcome measures | **Inclusion criteria:** IEGs expression as measured by mRNA or protein expression in one of the following brain regions: amygdala, hippocampus, hypothalamus, medial prefrontal cortex, nucleus accumbens, striatum; IEGs expression in other brain regions will be included only at a systematic review level  **Exclusion criteria:** brain regions not specified |  |
| 27. | Language restrictions | **Inclusion criteria:**  **Exclusion criteria:** None |  |
| 28. | Publication date restrictions | None |  |
| 29. | Other | The second hypothesis-confirming research question asks about the effects of multiple hits on the relationship between ELA and IEGs expression.  The following events are considered second hits:   - Stressful behavioral test performed previously (e.g., FST, fear conditions) - Footshocks - Chronic (mild) unpredictable stress - Chronic constant light - Chronic restraint - Chronic individual housing - Vaginal balloon distention - Cannula implementation, mock surgeries, blood sampling, isoflurane anaesthesia - Dams transported pregnant - Stress prone strain (BALB/C, wistar Kyoto, DBA)   The following events are not classified second hits:   - Intragastric saline - Saline injections - Vaginal smears - Daily handling by experimenter |  |
| 30. | Sort and prioritize your exclusion criteria per selection phase | **Titles and abstracts selection:**   1. Not primary publications. 2. Did not use mice/rats. 3. Not a model of early life adversity.   **Full text selection:**   4. Did not measure IEG products (mRNA or protein expression). 5. Not an acute stressor in adult life. 6. Animals fall into any of the exclusion criteria as specified in question 19. 7. Interventions fall into any of the exclusion criteria as specified in question 20. 8. Outcome measures fall into any of the exclusion criteria as specified in question 24. 9. Intervention specific to control group/experimental group, so that the groups differ by more than just early life adversity exposure. |  |
|  | Study characteristics to be extracted (for assessment of external validity, reporting quality) | | |
| 31. | Study ID (*e.g.* authors, year) | Study ID  Author  Abstract  Year  Journal |  |
| 32. | Study design characteristics (*e.g.* experimental groups, number of animals) | n Control (only differs by exposure to ELS)  n Experimental (ELS) |  |
| 33. | Animal model characteristics (*e.g.* species, gender, disease induction) | Species  Strain  Origin of the animals (own breeding, purchased pregnant, etc.)  Sex  Age at Experiment (Acute Stressor) |  |
| 34. | Intervention characteristics (*e.g.* intervention, timing, duration) | 1. **ELA model:** model, duration, start (related to age of animal), end (related to age of animal), litter size 2. **Acute stressor:** type, categorization, duration, intensity (if applicable), time (of day), time before death 3. **Multiple hits:** yes/no, type (if applicable) |  |
| 35. | Outcome measures | IEG (name; categorical)  Brain area (name; categorical)  Type (e.g., mRNA or protein; categorical)  Measure (e.g., percentage, fold-increase, optical density, counts; categorical) |  |
| 36. | Other (*e.g.* drop-outs) | / |  |
|  | Assessment risk of bias (internal validity) or study quality | | |
| 37. | Specify (a) the number of reviewers assessing the risk of bias/study quality in each study and (b) how discrepancies will be resolved | Risk of bias will be assessed by two independent researchers. Risk of bias is assessed following SYRCLE guidelines, and it will be distinguished between experimental and study bias. Discrepancies will be resolved by discussion between two experimenters. Should no conclusion be reached between two experimenters, a third researcher (expert in the field of early life adversity), will be consulted for a solution. |  |
| 38. | Define criteria to assess (a) the internal validity of included studies (*e.g.* selection, performance, detection and attrition bias) and/or (b) other study quality measures (*e.g.* reporting quality, power) | ☑By use of [SYRCLE's Risk of Bias tool^4^](http://www.biomedcentral.com/1471-2288/14/43/abstract)  □By use of SYRCLE’s Risk of Bias tool, adapted as follows:  □By use of [CAMARADES' study quality checklist, e.g ^22^](http://www.ncbi.nlm.nih.gov/pubmed/15060322)  □By use of CAMARADES' study quality checklist, adapted as follows:  □Other criteria, namely: |  |
|  | Collection of outcome data | | |
| 39. | For each outcome measure, define the type of data to be extracted (*e.g.* continuous/dichotomous, unit of measurement) | Mean Control (continuous)  Mean Experimental (continuous)  Standard Deviation Control (continuous)  Standard Deviation Experimental (continuous)  Reported direction of the effect (increase, decrease, non-significant; categorical)  Data will be extracted in form of a comparison between a control and an experimental group, which only differ in exposure to early-life adversity. The same animal (group) can be part of multiple comparisons. |  |
| 40. | Methods for data extraction/retrieval (e.g. first extraction from graphs using a digital screen ruler, then contacting authors) | 1. Extraction from numbers provided in the text (means, standard deviations, n).  2. Extraction from graphs.  3. Extraction from statistical analyses.  4. Contacting the authors. |  |
| 41. | Specify (a) the number of reviewers extracting data and (b) how discrepancies will be resolved | (a) One reviewer will complete data extraction, with a second reviewer checking random samples for agreement. Any numbers presented in the article or supplementary material will be extracted. If data is only presented graphically, then ‘WebPlotDigitizer’ will be used to extract data from graphs. If results of statistical analyses are given, these will be used to infer summary statistics. Two authors per publication will be contacted in case of missing data, followed by a reminder in case of no reply. Should authors not answer within two months, the comparisons will be reported as missing and will be excluded from analyses.  (b) Discrepancies will be resolved by discussion between two experimenters. Should no conclusion be reached between two experimenters, a third researcher (expert in the field of early life adversity), will be consulted for a solution. |  |
|  | Data analysis/synthesis | | |
| 42. | Specify (per outcome measure) how you are planning to combine/compare the data (*e.g.* descriptive summary, meta-analysis) | A quantitative synthesis is planned for results concerning the immediate early genes c-fos, arc, and egr1. Data will be split by sex, and the meta-analysis will be conducted for each dataset separately, since we consider males and females to be two different biological systems that should not be grouped together. |  |
| 43. | Specify (per outcome measure) how it will be decided whether a meta-analysis will be performed | The decision on which brain regions and acute stressors to include in the quantitative analysis will be made after study selection, with frequency being the determining factor. Remaining immediate early genes, brain regions and acute stressors, as well as the early life adversity model of handling, will be covered in a narrative/descriptive synthesis. |  |
|  | *If a meta-analysis seems feasible/sensible, specify (for each outcome measure):* | | |
| 44. | The effect measure to be used (*e.g.* mean difference, standardized mean difference, risk ratio, odds ratio) | The standardized mean difference Hedge’s g ( (mean(Control) - mean(Experimental)) / pooled SD ) will be used for all outcome measures. |  |
| 45. | The statistical model of analysis (*e.g.* random or fixed effects model) | 3-level mixed effect meta-analysis (in case of multiple outcomes from the same animals) otherwise random effects meta-analysis, with early life stress predicting IEG mRNA and protein expression. IEG identity, presence of second hits and brain region (if applicable) will be moderators in the model. |  |
| 46. | The statistical methods to assess heterogeneity (*e.g.* I^2^, Q) | Cochranes Q-test; I^2^ |  |
| 47. | Which study characteristics will be examined as potential source of heterogeneity (subgroup analysis) | Type of IEGs, ELA models, species, types of acute stressors, brain area and outcome measure (mRNA vs protein) will be used for subgroup analyses. These will be considered exploratory. |  |
| 48. | Any sensitivity analyses you propose to perform | Specified prior to the analysis, we will assess the influence of the following factors on the outcome measure:   1. Influential cases and outliers. 2. Blinded and randomized studies. 3. Risk of potential bias (bias will be assessed with the SYRCLE Risk of bias tool and an overall score will be used for sensitivity analysis). |  |
| 49. | Other details meta-analysis (*e.g.* correction for multiple testing, correction for multiple use of control group) | **Correction for multiple testing:** Bonferroni for family-wise comparisons will be applied for subgroup and exploratory analyses. Primary hypothesis-confirming research questions are considered separate families.  **Correction for multiple use of control group:** n (control) / n (comparison) | Change: Holm for p-values correction |
| 50. | The method for assessment of publication bias | If sufficient number of studies is achieved and a meta-analysis is conducted, the following methods will be applied:   1. Qualitative assessment of funnel plot 2. Egger’s regression, followed by test for funnel plot asymmetry 3. Begg’s test 4. Fail and save test 5. Trim and fill |  |

#### 1.2 Search String

**Pubmed:**

*Part 1 - Mice and rats:*

("rodentia"[Mesh] OR rodent*[tiab] OR "mus"[Tiab] OR "mice"[Mesh] OR "mice"[tiab] OR "mouse"[tiab] OR "rats"[Mesh] OR "rats"[tiab] OR "rat"[tiab])

*Part 2 – Postnatal early-life adversity:*

("maternal behavior"[MeSh] OR "maternal care”[tiab] OR "early life stress"[tiab] OR "ELS"[tiab] OR "early life adversity"[tiab] OR "early life adversities"[tiab] OR "ELA"[tiab] OR "early life manipulation"[tiab] OR "early life manipulations"[tiab] OR "early adverse experience"[tiab] OR "early adverse experiences"[tiab] OR "early adversed experience"[tiab] OR "early adversed experiences"[tiab] OR "perinatal stress"[tiab] OR "perinatal adversity"[tiab] OR "perinatal adversities"[tiab] OR "perinatal manipulation"[tiab] OR "perinatal manipulations"[tiab] OR "perinatal adverse experience"[tiab] OR "perinatal adverse experiences"[tiab] OR "perinatal adversed experience"[tiab] OR "perinatal adversed experiences"[tiab] OR "postnatal stress"[tiab] OR "postnatal adversity"[tiab] OR "postnatal adversities"[tiab] OR "postnatal manipulation"[tiab] OR "postnatal manipulations"[tiab] OR "postnatal adverse experience"[tiab] OR "postnatal adverse experiences"[tiab] OR "postnatal adversed experience"[tiab] OR "postnatal adversed experiences"[tiab] OR "neonatal stress"[tiab] OR "neonatal adversity"[tiab] OR "neonatal adversities"[tiab] OR "neonatal manipulation"[tiab] OR "neonatal manipulations"[tiab] OR "neonatal adverse experience"[tiab] OR "neonatal adverse experiences"[tiab] OR "neonatal adversed experience"[tiab] OR "neonatal adversed experiences"[tiab] OR "Maternal Deprivation"[Mesh] OR "maternal deprivation”[tiab]OR "maternal separation”[tiab] OR "limited bedding”[tiab] OR "limited nesting”[tiab] OR "limited material”[tiab] OR "limited bedding/nesting"[tiab] OR "limited bedding-and-nesting"[tiab] OR "limited nesting/bedding"[tiab] OR "limited nesting-and-bedding"[tiab] OR "early life isolation"[tiab] OR "perinatal isolation"[tiab] OR "postnatal isolation"[tiab] OR "neonatal isolation"[tiab] OR "licking and grooming"[tiab] OR "licking-and-grooming"[tiab] OR "licking/grooming"[tiab] OR "early handling"[tiab] OR "early life handling"[tiab] OR "perinatal handling"[tiab] OR "postnatal handling"[tiab] OR "neonatal handling"[tiab])

**Embase search string**

*Part 1 – Mice and rats:*

(rodent*:ab,ti OR mus:ab,ti OR mouse:ab,ti OR mice:ab,ti OR rat:ab,ti OR rats:ab,ti)

*Part 2 – Postnatal early-life adversity:*

('maternal behavior':ab,ti OR 'maternal care':ab,ti OR 'early life stress':ab,ti OR 'els':ab,ti OR 'early life adversity':ab,ti OR 'early life adversities':ab,ti OR 'ela':ab,ti OR 'early life manipulation':ab,ti OR 'early life manipulations':ab,ti OR 'early adverse experience':ab,ti OR 'early adverse experiences':ab,ti OR 'early adversed experience':ab,ti OR 'early adversed experiences':ab,ti OR 'perinatal stress':ab,ti OR 'perinatal adversity':ab,ti OR 'perinatal adversities':ab,ti OR 'perinatal manipulation':ab,ti OR 'perinatal manipulations':ab,ti OR 'perinatal adverse experience':ab,ti OR 'perinatal adverse experiences':ab,ti OR 'perinatal adversed experience':ab,ti OR 'perinatal adversed experiences':ab,ti OR 'postnatal stress':ab,ti OR 'postnatal adversity':ab,ti OR 'postnatal adversities':ab,ti OR 'postnatal manipulation':ab,ti OR 'postnatal manipulations':ab,ti OR 'postnatal adverse experience':ab,ti OR 'postnatal adverse experiences':ab,ti OR 'postnatal adversed experience':ab,ti OR 'postnatal adversed experiences':ab,ti OR 'neonatal stress':ab,ti OR 'neonatal adversity':ab,ti OR 'neonatal adversities':ab,ti OR 'neonatal manipulation':ab,ti OR 'neonatal manipulations':ab,ti OR 'neonatal adverse experience':ab,ti OR 'neonatal adverse experiences':ab,ti OR 'neonatal adversed experience':ab,ti OR 'neonatal adversed experiences':ab,ti OR 'maternal deprivation':ab,ti OR 'maternal separation':ab,ti OR 'limited bedding':ab,ti OR 'limited nesting':ab,ti OR 'limited material':ab,ti OR 'limited bedding/nesting':ab,ti OR 'limited bedding-and-nesting':ab,ti OR 'limited nesting/bedding':ab,ti OR 'limited nesting-and-bedding':ab,ti OR 'early life isolation':ab,ti OR 'perinatal isolation':ab,ti OR 'postnatal isolation':ab,ti OR 'neonatal isolation':ab,ti OR 'licking and grooming':ab,ti OR 'licking-and-grooming':ab,ti OR 'licking/grooming':ab,ti OR 'early handling':ab,ti OR 'early life handling':ab,ti OR 'perinatal handling':ab,ti OR 'postnatal handling':ab,ti OR 'neonatal handling':ab,ti)

#### 1.3 Extracted Variables

To increase subjectivity during data extraction, variables to be extracted were determined *a priori.* The spreadsheet containing all extracted variables and variable coding is available at <https://osf.io/qkyvd/>.

| Entity | Variables |
| --- | --- |
| Publication | title; authors; year; journal |
| Animal | species; strain; origin (e.g., breeding, dams purchased pregnant); sex |
| Model | model (type, timing, cage (novel cage or home cage); light/dark phase; repetition (e.g., once, twice, predictable, unpredictable)); cross fostering; culling; sex ratio; litter size |
| Multiple Hits | other life experiences; housing in adulthood |
| Testing | age at testing; acute stressor (type, duration and novelty); time until perfusion; estrous cycle phase (females only) |
| Outcome | IEG name and product; measurement (technique, unit of recording (e.g., counts, expression, optical density) and unit of comparison (e.g., raw data, fold change, averages across slices)); brain area and hemisphere |
| Data | mean, variance and n of control and experimental groups; significant effect |

#### 1.4 Variables’ grouping

A) Brain areas as named in publications and as grouped for the analysis.

| **Grouped for analysis** | **Named in publications** |
| --- | --- |
| **Amygdala** | Central amygdala; Medial amygdala; Basolateral nucleus; Amygdala central nucleus; basolateral amygdala; Lateral amygdala |
| **Hippocampus** | Dorsal CA1; Dorsal CA2; Dorsal CA3; Dorsal dentate gyrus; Ventral CA1; Ventral CA2; Ventral CA3; Ventral dentate gyrus; CA1 subregion of the hippocampus; Dentate gyrus; Central CA3; CA1 region; CA1; CA2; CA3; Hippocampus |
| **Hypothalamus** | PVN; mpPVN; mgPVN; lpPVN; dpPVN; Medial mammillary nucleus; medial parvocellular portion of the PVN; paraventricular nucleus of the HAT; Paraventricular nucleus; ventromedial hypothalamic nucleus; anterior hypothalamus; lateral hypothalamus; dorsolmedial hypothalamus |
| **Prefrontal cx** | ACC; Cingulate cortex; mPFC; caudal cingulate cortex; rostral cingulate cortex; infralimbic cortex; prelimbic cortex; anterior cingulate cortex; prefrontal cortex; Cingulate cortex; lateral orbital frontal cortex; medial orbital frontal cortex; ventral orbital frontal cortex; medial prefrontal cortex |
| **Thalamus** | CM; PV; VPL; Anterodorsal thalamic nuclei; central medial thalamic nucleus; anteroventral thalamus; anteromedial thalamus |
| **Other*** | dorsal striatum; Barrel cortex; Piriform cortex; Lateral septum; Caudate putamen; DRN; Pontine region; Cerebellum; vBNST; Nucleus accumbens; ventrolateral periaqueductal gray; dorsolateral periaqueductal gray; DRD; DRV; DRVL; DRI; Retrosplenial cortex; non-preganglionic Edinger-Westphal nucleus; dorsal raphe nucleus; forebrain neocortical tissue; Nacc; VTA; medial orbital frontal cortex; ventral orbital frontal cortex; lateral orbital frontal cortex; insular cortex; dorsolateral striatum; dorsomedial striatum; nucleus accumbens shell; Cortex; Striatum; periaqueductal gray; bed nuclues of the stria terminalis; ventral subiculum; dorsal lateral septum; ventral lateral septum; medial septum; dorsal periaqueductal gray; inferior colliculus; locus coereleus; lateral septum; nucleus accumbens; ventral pallidum |

* ‘Other’ brain areas have not been included in the analysis.

B) Categorization of acute stress in mild and severe.

| **Intensity** | **Type** |
| --- | --- |
| **Mild** | EPM; OFT; DLB; Competition; Reexposure FC context (no shock); Social Defeat; NE; IGT; Three chamber test; Social interaction after 1d of social isolation |
| **Severe** | CRD; RS; FS in inhibitory avoidance task; FST; Shock in shock-probe burial task; MWM |

Of note, stressors with a strong memory, social or reward component were assessed on a systematic review level only.

### 2. Supplementary Results

#### 2.1 Bias assessment

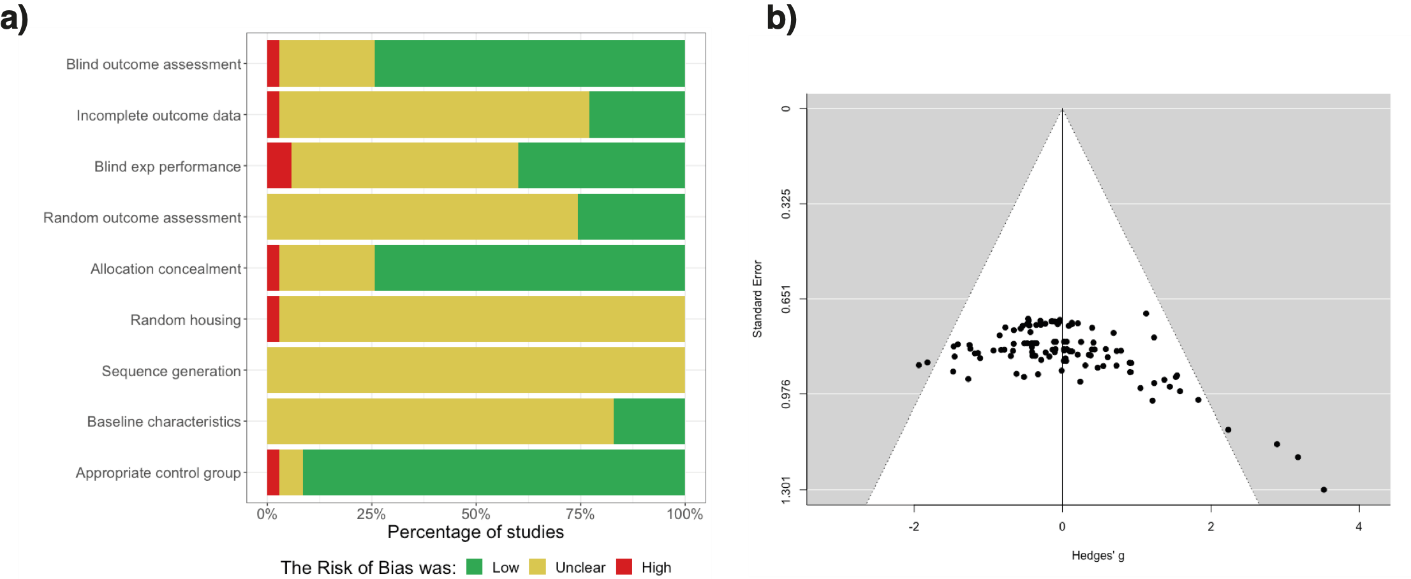

**Bias assessment.** A) Risk of bias assessment according to SYRCLE’s risk of bias tool. B) Funnel plot for publication bias.

#### 2.2 Systematic review

#### 2.2.1 cFos in female rodents.

Given fundamental biological differences between males and females [1], we *a priori* chose to evaluate female cFos data separately from males’. Only ten publications reported on cFos expression in female rodents (*n_comp_* = 77). The majority of these studies found no significant differences between cFos levels of ELA versus controls at rest or after an acute stress challenge (n_comp_ = 55; [2–6]).

Only five studies performed the same experiments in both male and female rodents. Among these, Desbonnet et al. [5], Gaszner et al. [6] and Renard et al. [2] reported the same null effects for both male and females. In contrast, James et al. [3] and Genest et al. [4] found no significant ELA effects on cFos levels in females, while they did report significant differences in males under the same conditions. These sexually dimorphic results could have methodological origins, such as male-focused behavioral paradigms (for ELA or acute stress) or could reflect true biological differences between the sexes [1, 7].

The remaining five studies investigated exclusively females, and all reported at least one significant difference between ELA and control rodents. Auth et al. [8] found significantly increased cFos levels in female mice at rest, but not after acute stress exposure. Interestingly, across two independent baseline cohorts, increased cFos was observed once in the dorso-lateral periaqueductal gray and once in the lateral amygdala, suggesting that the effects do not easily replicate within the same lab. Similarly, Rivarola and colleagues [9, 10] observed an increase in cFos levels in the anterior-dorsal thalamic nucleus of animals with a history of multiple hits in a first [9] but not a second publication [10].

Finally, O’Leary et al. [11] reported decreased cFos levels in the dorsal dentate gyrus and ventral CA3 of female ELA mice after restraint stress, but not in other hippocampal, hypothalamic, prefrontal cortical or amygdalar areas. Banqueri et al. [12] demonstrated differential directionality of effects after the Morris water maze, with ELA females showing increased cFos levels in hippocampal structures, and decreased expression in prefrontal areas. All in all, ELA effects on cFos in females appeared limited. Whether the results are truly sexually dysmorphic remains to be elucidated.

#### 2.2.2 cFos and other brain areas

Five studies investigated the effect of ELA on cFos expression in brain areas of male rodents other than those reviewed in the meta-analysis, including the striatum, sensory cortices, hindbrain nuclei and the cerebellum of male rodents. Out of 24 comparisons, 16% displayed a significant difference between ELA and control animals (*n_comp_* = 4) at systematic review level. Troakes et al. [13] showed that cFos levels of ELA males are significantly decreased in the piriform cortex in comparison to controls after acute exposure to a mild stressor, but not at rest. Early research indicated that cFos levels in the piriform cortex are highly responsive to acute stressors, and its role in the sensory integration of olfactory stimuli suggests that the reduced cFos expression could correspond to decreased information processing abilities under stressful [14, 15].

In addition, Menard et al. [16] found decreased cFos expression in ELA males in the lateral septal complex and the ventral subiculum after performing a shock-probe burial task, but not in other striatal areas or hindbrain nuclei. Given that the lateral septal complex relays reward and fear information for contextualization of the experience, the decreased cFos expression here potentially presents a task-specific effect related to spatial mapping of the buried probe [17]. However, Shin et al. [18] report upregulation of cFos after ELA in the lateral septal complex as well as the ventral tegmental area in a social interaction task, suggesting a broader task-specific involvement of striatal areas.

Finally, neither Clarke et al. [19] nor Desbonnet et al. [5] could find significant differences between ELA males and controls in the bed nuclei of the stria terminalis, neither at rest nor after acute stress, suggesting that cFos expression in this area is not or only minimally changed after ELA. All in all, these results suggest that areas with task-specific effects are worth exploring, and that cortical areas involved in sensory processing and information integration potentially display altered transcriptional activity as well. Yet, considering that the most frequently areas under investigation are also those areas considered to be sensitive to the effects of stress, it is likely that the main results of interest are covered by the meta-analytic outcomes.

#### 2.2.3 cFos and alternative behavioral paradigms.

Acute stressors that included a strong memory, reward or social component were excluded from the meta-analysis. They involve cognitive processes other than the response to stress, which recruit brain-areas depending on the task requirements.

Daskalakis et al. [20] investigated cFos expression in rats placed back into a fearful context after a fear-conditioning paradigm, thereby probing memory processes in addition to stress-related functions. cFos expression in the medial amygdala and basolateral amygdala was increased in rats placed into a novel cage during the *maternal separation* (MS) procedure, while an increase was only observed in the medial amygdala in MS animals that remained in the home cage [20]. This study highlights how that choices of study characteristics (i.e., home cage vs novel cage) can influence the outcome investigated.

Two studies further investigated the effects of ELA on cFos expression after exposure to a rodent version of the Iowa Gambling Task [21, 22]. This task depends not only on spatial memory, but also contains a strong reward component [22]. In 2012, van Hasselt et al. [21] correlated percentage of licking and grooming with cFos expression in a wide range of brain areas in male and female rats and found a negative correlation in the shell of the nucleus accumbens and the agranular insular cortex when sex was pooled. However, using the same task, MS did not alter cFos expression in these areas in the 2017 study, but rather decreased cFos expression in the right CA1, right CA3, left infralimbic area and left agranular insula [22]. While inconsistent, these studies highlight that reward-based processes also likely result in differential activation of IEGs after ELA exposure, thus, warranting further investigations in the future.

Under several social paradigms, no differences between ELA and control animals were observed in medial PFC areas [18, 23, 24], the central amygdala [23], the dorsal raphe nucleus [25], or striatal and hypothalamic areas [18]. On the other hand, Benner et al. [23] observed an increase in the basolateral amygdala and a decrease in CA1 of cFos expression in ELA mice compared to controls after 40-days social competition task. A possible explanation is that the differences observed in the study by Benner and colleagues are due to the memory component, rather than the stress/social component of the task. In addition, Shin et al. [18] observed an increase in cFos expression in the lateral septal complex and the ventral tegmental area after social interaction in mice previously exposed to social isolation, suggesting that multiple adverse experiences may be required to observe altered IEG expression after ELA in social tasks. Overall, social behaviors in isolation seem less inducive of activity-regulated transcription than the above-discussed reward-based and memory-based paradigms.

#### 2.2.4 ELA and IEG other than cFos

*Arc* is a post-synaptic protein, which plays an essential role in regulating the homeostatic scaling of AMPA receptors, thereby directly modifying plasticity at the synapse [26]. *Arc* expression has been investigated in five publications under varying conditions in male and female mice and rats. While two publications did not find any alterations in the mPFC, hippocampal, or amygdaloid areas at rest or after acute stress [23, 27], another study reported a significant decrease in CA1, CA3 and dentate gyrus *Arc* levels in male ELA animals at rest [28]. Interestingly, animals in this study were exposed to maternal separation for one week longer (PND 1-21) than animals in the studies reporting no significant alterations, suggesting that the duration of the ELA experience could be essential in causing long-term effects on *Arc* expression. It is noteworthy that a decrease in *Arc* expression results in increased synaptic plasticity [26], thus, following in line with the findings of increased cFos expression at rest in the male meta-analysis.

In contrast, McGregor et al. [29] found increased *Arc* expression at rest in the dorsal striatum of male rats with and without a history of second hits. As this publication is the only one reporting on IEG levels in the dorsal striatum, it is unclear whether the finding is a result of the study design or presents a genuine area specific IEG response. Rincel et al. [30] suggest that ELA effects on Arc expression are sex-specific, showing evidence that ELA leads to decreased Arc expression in the mPFC of male mice, but to increased Arc levels in the mPFC of female mice. These contradictory findings could be a strain-specific, as C3H/HeNRj mice were used [30]. All in all, reported Arc levels appear to be in coherence with cFos effects on synaptic plasticity at rest, and thereby further support the notion of at rest sensitization of activity-regulated transcription.

Early-growth response (Egr) proteins are a family of transcription factors with a zinc-finger motif, which allows all *Egr* factors to connect to identical DNA binding sites [31]. We identified three studies investigating Egr expression after ELA exposure at rest; one investigated Egr-1 [32], another investigated Egr-4 only [30], and one other investigated Egr-2 and Egr-4 [29].

*Egr-1* mRNA expression was decreased in the cortex, but only in Balb/c and not C57Bl/6 male mice [32]. This is in line with the general notion of Balb/c mice as a stress-sensitive strain [33]. In contrast, McGregor et al. [29] report increases in Egr-2 and Egr-4 expression in the dorsal striatum, with Egr-2 levels only significantly increased in animals experiencing a second hit during adolescence. Finally, Rincel et al. [30] highlight that ELA alters Egr-4 expression in a sex-specific manner in the mPFC of mice, observing an downregulation in males but an upregulation in females.

Since it is expected that proteins of the Egr-family behave similarly [31], the discrepancy between findings are likely the result of differences in study design, such as the brain areas investigated. Considering that IEGs of the Egr*-*family, and in particular *Egr-1,* have been shown to be associated with the development and treatment of those psychiatric disorders, which individuals with a history of early life stress are more likely to develop, Egr*-*family proteins are an understudied, yet important candidate for investigating activity-regulated transcriptional alterations after ELA in the future [34].

*FosB* is an IEG of the *Fos* family, and - similarly to *cFos* - if binds to members of the *Jun* family to form the AP1 transcription factor [35]. Of particular interest in stress research is its isoform *ΔFosB*, whose extended half-life makes *ΔFosB* an exceptional marker for chronic stress [35].

Three publications reporting on the expression of *ΔFosB* at rest in ELA and control animals were identified. Kim et al. [36] reported a reduction of *ΔFosB* expression in the nucleus accumbens of ELA females in comparison to controls, whereas Wang et al. [37] report elevated *ΔFosB* levels in the mPFC of ELA rats of unspecified sex, pointing towards opposite effects of ELA in these two areas. Interestingly, and in line with these findings, previous results suggest that overexpression of *ΔFosB* in the nucleus accumbens accompanied by reduced expression of *ΔFosB* in mPFC promote a phenotype resilient to the effects of chronic stress [38, 39]. It should still be highlighted, that Lippmann et al. [40] found no significant alterations in either of these areas in male rodents, neither induced by maternal separation nor by handling. Due to the low number of studies investigating *ΔFosB*, we cannot conclude whether these null findings are attributable to sex or a result of study design heterogeneity. Yet the outlined potential of a more stable IEG in researching chronic alterations in transcriptional activity emphasizes the relevance of investigating ELA modifications on *ΔFosB* expression.
